# Electron Microscopic Reconstruction of Neural Circuitry in the Cochlea

**DOI:** 10.1101/2020.01.04.894816

**Authors:** Yunfeng Hua, Xu Ding, Haoyu Wang, Fangfang Wang, Yunge Gao, Yan Lu, Tobias Moser, Hao Wu

**Affiliations:** Department of Otolaryngology-Head and Neck Surgery, Shanghai Ninth People’s Hospital, China; Ear Institute, Shanghai Jiao Tong University School of Medicine, China; Shanghai Key Laboratory of Translational Medicine on Ear and Nose Diseases, China; Shanghai Institute of Precision Medicine, Shanghai Ninth People’s Hospital, China; Department of Connectomics, Max-Planck-Institute for Brain Research, Germany; Institute for Auditory Neuroscience, University Medical Center Göttingen, Germany; Auditory Neuroscience Group, Max Planck Institute of Experimental Medicine, Germany; Multiscale Bioimaging Cluster of Excellence (MBExC), University of Göttingen, Germany

**Author notes:** These authors contributed equally to this work (Yunfeng Hua, Xu Ding, Haoyu Wang, Fangfang Wang). Correspondence (H.W.), (T.M.).

## Abstract

Recent studies have revealed great diversity in the structure, function and efferent innervation of afferent synaptic connections between the cochlear inner hair cells (IHCs) and spiral ganglion neurons (SGNs), which likely enables audition to process a wide range of sound pressures. By performing an extensive electron microscopic reconstruction of the neural circuitry in the mature mouse organ of Corti, we demonstrate that afferent SGN-dendrites differ in strength and composition of efferent innervation in a manner dependent on their afferent synaptic connectivity with IHCs. SGNs that sample glutamate release from several presynaptic ribbons receive more efferent innervation from lateral olivocochlear projections than those driven by a single ribbon. Next to the prevailing unbranched SGN-dendrites, we found branched SGN-dendrites that can contact several ribbons of 1-2 IHCs. Unexpectedly, medial olivocochlear neurons provide efferent innervation of SGN-dendrites, preferring those contacting single-ribbon, pillar-side synapses. We propose a fine-tuning of afferent and efferent SGN-innervation.

## Introduction

Acoustic information is encoded into neural signals in the cochlea, the mammalian hearing organ in the inner ear. Sound encoding by afferent spiral ganglion neurons (SGNs) faithfully preserves signal features such as frequency, intensity, and timing (for reviews see ref. 1–3). It also is tightly modulated by efferent projections of the central nervous system to enable selective attention and to aid signal detection in noisy background, sound source localization as well as ear protection against acoustic trauma (see reviews^4,5^). Sound encoding occurs at the afferent connections between inner hair cells (IHCs) and postsynaptic type-1 SGNs (^1^SGNs) that constitute 95% of all SGNs^6,7^. These so-called ribbon synapses exhibit glutamate-mediated neurotransmission at rates of hundreds per second and with sub-millisecond precision^1,8^.

In the mouse cochlea, each IHC is contacted by 10-30 ^1^SGNs and it is thought that each IHC active zone drives firing in a single unbranched peripheral SGN-neurite (also referred to as SGN-dendrite). These afferent connections exhibit great diversity in synaptic and ^1^SGN dendritic morphology^9–15^, physiological properties^9,12,14,16^, and molecular ^1^SGN profile^17–19^. Across different animal species, ^1^SGN can be classified into three functional subtypes, namely low, medium, and high spontaneous rate (SR) ^1^SGNs that also differ in the thresholds and dynamic range of sound encoding^7,16,20–22^. In the cat^6^, classical back-tracing experiments linked function to morphology showing that low SR ^1^SGNs have thinner dendrites with fewer mitochondria than high SR ^1^SGNs. Low SR ^1^SGNs preferentially innervate the modiolar (neural) side of the IHC, where they face larger and more complex presynaptic active zones (AZs)^6,23–25^. Larger and more complex AZs at the modiolar side of IHCs were also found in mouse^11–13^, guinea pig^26,27^, and gerbil^28^, suggesting that some morphological corollaries of functional ^1^SGN diversity may be shared across species. Modiolar AZs, in general, contain larger and/or multiple ribbons as well as more membrane-proximal SVs^13,23^ and Ca^2+^ channels that need stronger depolarization for activation than at the pillar (abneural) face^12^. What might appear as a peculiar biological variance at first glance, is becoming increasingly recognized as a fascinating parallel information processing mechanism by which the auditory system copes with encoding over a broad range of sound pressures downstream of cochlear micromechanics. Specifically, emerging evidence indicates that the full sound intensity information contained in the IHC receptor potential is fractionated into subpopulations of ^1^SGNs that collectively encode the entire audible range of ^1^SGNs (recent reviews in ref. 1,29,30). Besides, afferent cochlear signaling is dynamically modulated by efferent projections emanating from the medial and lateral superior olivary complex of the auditory brainstem^31^. The efferent projections of the medial olivocochlear component (MOC) form cholinergic synapses onto outer hair cells (OHCs) and suppress their electromotility, thereby attenuating cochlear amplification^32–34^. The lateral olivocochlear component (LOC) directly modulates ^1^SGNs via axodendritic synapses near the afferent ribbon synapses (for reviews see ref. 5,35). Efferent innervation seems more pronounced for ^1^SGNs contacting IHCs at the modiolar face and de-efferentation interferes with the modiolar-pillar gradient of ribbon size^36^. In rodents, the LOC fibers express a variety of neurotransmitters including dopamine (DA), acetylcholine (ACh) and ϒ-amino-butyric acid (GABA) among others (for reviews see ref. 37,38). LOC control of ^1^SGNs is thought to serve interaural matching^39,40^ as well as ^1^SGN gain control^41^.

Despite major efforts on structural investigation of cochlear afferent and efferent innervation using light microscopy^42–45^ and electron microscopy (EM)^15,23,24,46–53^, a comprehensive diagram of the neural circuitry in the organ of Corti remains to be established. Here we report the complete reconstruction of two large EM volumes acquired from the organ of Corti of the mid-frequency cochlear region of mature CBA mice. We employed serial block-face EM (SBEM) imaging^54^ which offered the spatial resolution required for both comprehensive connectomic analysis and structural studies of afferent and efferent synapses. This revealed a substantial fraction of ^1^SGNs (12 %) with terminal bifurcation, which results in input of two or more AZs from one or two IHCs. Unexpectedly, we found robust MOC innervation of ^1^SGN dendrites preferentially on the pillar IHC side. Efferent innervation, primarily by LOCs, was stronger for ^1^SGN dendrites receiving input from multiple ribbons.

## Results

To perform a comprehensive structural analysis of the mammalian cochlear circuit, we first acquired a SBEM dataset from the left cochlea of a 7-week-old wildtype female mouse (**Fig. 1 a** & **b**). We employed CBA mice that are considered as the gold standard for analyzing normal mouse hearing as they, in contrast to C57BL/6 mice, do not show rapid age-related hearing loss^55^. The field of scanning was centered on the organ of Corti of the cochlear mid-turn and encompassed 15 IHCs and 45 OHCs. 2500 consecutive image slices were collected, resulting in a 3D-aligned volume of (xyz) 262 × 194 × 100 μm^3^ at 11 × 11 × 40 nm^3^ voxel size (**Suppl. video 1** & **2**). Dense reconstruction of the inner spiral bundle (ISB) beneath the 15 IHCs (**Fig. 1 c**) was done by manual annotation using an open-source visualization and annotation software called webKNOSSOS^56^. Despite 40 nm cutting thickness, quantification using a redundant-skeleton consensus procedure (RESCOP)^57^ suggests a tracing quality comparable to the published SBEM cortex datasets of 30 nm cutting thickness (**Suppl. Figure 1**). Tracing revealed a total of 234 ^1^SGN-dendrites, 32 LOC as well as 39 MOC fibers, which contribute 47.2 %, 36.5 %, and 13.0 % to a total circuit path length of 16.97 mm in the ISB region (**Fig. 1 d** & **e**). SGN-dendrites were identified as fibers contacting ribbon-type AZs of IHCs (dendrites emanating from ^1^SGNs) or OHCs (dendrites emanating from ^2^SGNs crossing the tunnel of Corti). Efferent fibers were defined as LOCs if they exclusively synapsed onto ^1^SGNs and IHCs (not further analyzed in this study) and as MOCs if they crossed the tunnel of Corti and also contacted the OHCs. All MOCs were thick compared to LOCs and ^2^SGNs and formed presynaptic terminals onto OHCs. A total of 1517 efferent synapses onto ^1^SGNs were annotated in the dataset. A second dataset (26 IHCs, 366 ^1^SGN-dendrites, **Fig. 1 f** & **g**) was acquired from an 8-week-old female mouse and analyzed after completing the analysis of the first EM volume.

**Figure 1.**
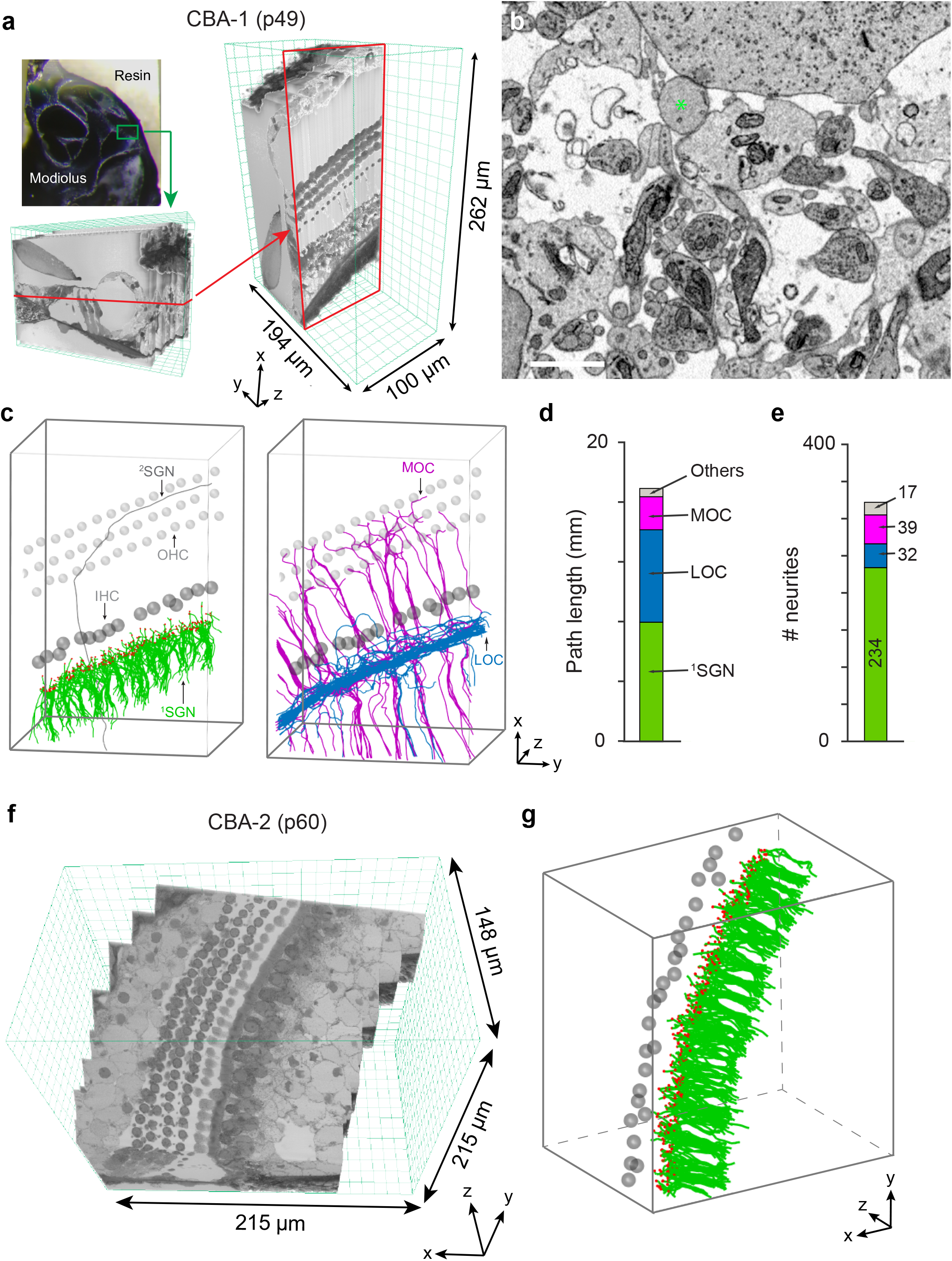
SBEM imaging and dense reconstruction of the mouse organ of Corti. **(a)** Left: location of the SBEM dataset (green box) in the resin-embedded cochlea. Right: dimension of the dataset. Red box: virtual cross-section at the position as indicated in (left). **(b)** High-resolution example image of neurites beneath IHCs and a representative ^1^SGN bouton was indicated by asterisk (green). Scale bar 2 μm**. (c)** Reconstruction of all 322 neurites in the dataset. Left: skeletons of all radial ^1^SGN-dendrites (green) and one representative ^2^SGN-dendrite with a characteristic turn towards the cochlear base (grey). Right: skeletons of all LOC fibers (blue) and MOC fibers (magenta) with main trunks extending into habenula perforatae and, for MOC, tunnel-crossing fibers towards the OHC region. Small grey spheres represent OHCs and large dark spheres represent IHCs. Scale bar 20 μm. **(d, e)** Quantification of circuit components. All 234 ^1^SGN-dendrites (green), 32 LOC fibers (blue) and 39 MOC fibers (magenta) contribute 47.2 % (8.01 mm), 36.46 % (6.19 mm) and 13.05 % (2.21 mm) of circuit path length (total: 16.97 mm) in the ISB region, respectively. Others include e.g. ^2^SGN-dendrites. **(f)** Snapshot of the second SBEM dataset. **(g)** Reconstruction of all 366 ^1^SGN-dendrites (green), all 468 ribbons (red) and 26 IHC nuclei (grey) of the second SBEM dataset. Scale bar 20 μm.

### Ribbon synapse heterogeneity in IHCs

Volume EM techniques have been recently employed to investigate ribbon synapse morphology in the apical cochlea of C57BL/6 mice at different developmental stages^13^. Our SBEM datasets now provide detailed structural insight into the mature, most sound sensitive, mid-cochlear region of the organ of Corti^10,16,42^. The high spatial resolution and large volume of the dataset allowed for precise structural quantification of ribbons as well as the postsynaptic ^1^SGN-dendrites (**Fig. 2 a** & **b**). From 15 reconstructed IHCs, a total of 308 ribbons were identified and volume traced by human annotators. Ribbon positions within IHCs were determined in the plane perpendicular to the IHC habenular-cuticular axis (same as described in ref. ^36^). The ribbon volume tended to be smaller in the 26 IHCs of the 2^nd^ animal which did not reach significance for the mean ribbon volume (0.0136 ± 0.0056, n = 308 for the 1^st^ animal vs. 0.0128 ± 0.0069, n = 468 for the 2^nd^ animal, p = 0.0932) but was significant for the ribbons on the pillar IHC side (**Fig. 2 c**, 0.0117 ± 0.0049 μm^3^, n = 142 vs. 0.0098 ± 0.0043, n = 192, p = 0.0183). We consider this to represent biological variance as technical reasons are unlikely (no indication for shrinkage based on other structures such as mitochondria or axon calibers for the 2^nd^ animal, data not shown).

**Figure 2.**
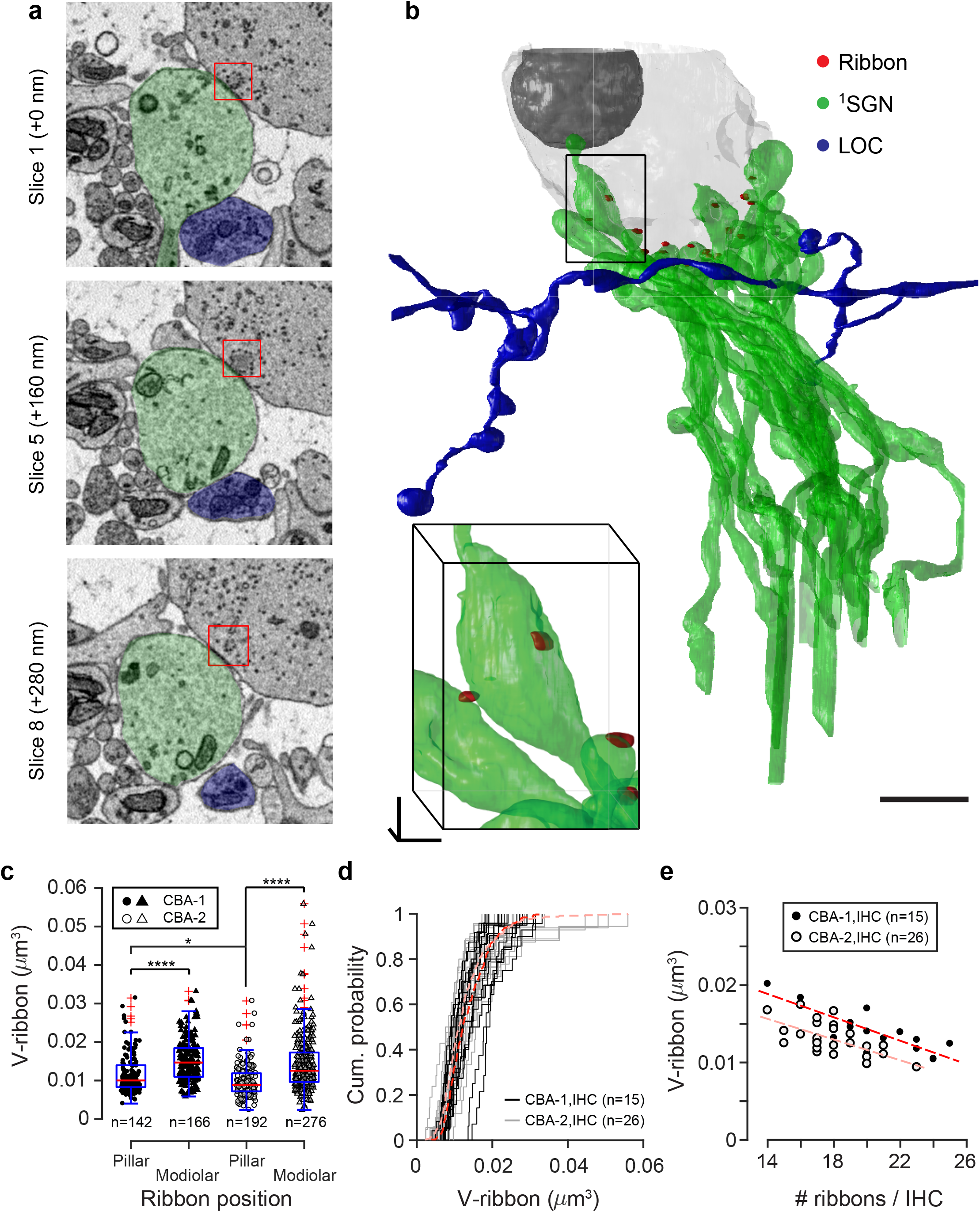
Quantification of synaptic ribbons in the IHCs. **(a)** Consecutive slices showing a representative ribbon synapse (red box) and the postsynaptic ^1^SGN bouton (green) co-innervated by an efferent terminal (blue). Scale bar 1 μm. **(b)** Volume reconstruction of an exemplary IHC with all ribbon synapses (red) and ^1^SGNs (green), and a representative LOC fiber (blue). Scale bar 5 μm. IHC location as indicated in Suppl. Figure 1. **(c)** Boxplot of measured ribbon sizes. All identified ribbons (308 in the 1^st^ animal and 468 in the 2^nd^ animal) were classified as two classes according to their locations (pillar or modiolar face of the IHCs). Exceptions are 12 (CBA-1) and 24 (CBA-2) cases with ribbons located at the bottom of IHCs that were grouped to the modiolar ribbons. Note that ribbons at the IHC pillar face (black dots and triangles, 0.0117 ± 0.0049 μm^3^ and 0.0098 ± 0.0043 μm^3^) were significantly smaller than their counterparts at the IHC modiolar face (circles and open triangles, 0.0153 ± 0.0055 μm^3^ and 0.0145 ± 0.0074 μm^3^). (Two-sample t-test, ****p < 0.0001). **(d)** Cumulative probability distribution of ribbon sizes from all 15 IHCs in the CBA-1 (red dash line) and 26 IHCs in the CBA-2 (light red dash line), as well as individual IHCs (CBA-1: black lines; CBA-2: grey lines). Note the remarkable cell-to-cell variability between IHCs. **(e)** Negative correlation between number and mean size of ribbons in individual IHCs (CBA-1: back dots; CBA-2: cycles). Linear fit: CBA-1 (red dash line, adjusted-R^2^ = 0.67) and CBA-2 (light red dash line, adjusted-R^2^ = 0.41).

In agreement with previous quantifications using light microscopy^11,12^, a prominent modiolar-pillar size gradient of ribbon sizes was observed in both animals (**Fig. 2 c**). On average, ribbons designated to the modiolar side of the IHCs were 40 % larger than those at the pillar side (in the 1^st^ animal, 0.0153 ± 0.0055 vs. 0.0117 ± 0.0049 μm^3^, p < 0.0001; in the 2^nd^ animal, 0.0147 ± 0.0074 vs. 0.0098 ± 0.0043 μm^3^, p < 0.0001). In addition to the heterogeneity of ribbon size within IHCs, we found differences in the distributions of ribbon abundance and size between IHCs, even in neighbors (**Fig. 2 d** & **e**), which have not yet been reported to our knowledge. In some extreme cases, individual IHCs can exclusively contain large or small ribbons, which we, different from^36^, found to be unrelated to the characteristic staggering arrangement of IHCs in the cochlear mid-turn. The amount of cellular ribbon material might be conserved as the mean size of ribbons in an IHC negatively correlates with its total number of ribbons (**Fig. 2 e**).

### Quantification of afferent and efferent inputs on ^1^SGN-dendrites

Besides receiving glutamatergic input from ribbon-type AZs of IHCs, ^1^SGN-dendrites are the primary postsynaptic target of efferent LOC projections. A prior study of the effects of lesioning the LOC projections indicated that innervation by LOCs is required for maintaining the modiolar-pillar ribbon size gradient^36^. Here, we mapped the efferent innervation on ^1^SGN-dendrites to relate it to the position of afferent ^1^SGN-boutons on IHCs and to the morphology of the corresponding IHC AZ. Reconstruction of ^1^SGN-dendrites together with their complete afferent and efferent inputs in the ISB region revealed several unexpected findings. First, we found that three morphologically distinct subpopulations of ^1^SGNs coexist in the mature cochlea (**Fig. 3 a**). Out of in total 234 ^1^SGNs in the 1^st^ animal, 170 (72.6 %) were identified as unbranched ^1^SGNs receiving input from an IHC AZs holding a single ribbon (“single-ribbon variant”, **Fig. 3 a1**), 35 (15.0 %) as unbranched ^1^SGNs but input from multiple ribbons (“unbranched multi-ribbon”), and strikingly 29 (12.4 %) as branched or bifurcated ^1^SGN-dendrites with input from multiple ribbons of one or two IHCs (“branched multi-ribbon”, **Fig. 3 a2** & **a3** and **Suppl. Figure 2**). Analysis performed in the 2^nd^ animal corroborated these results: this and further classification of ^1^SGN-dendrites of both animals are summarized in **Table 1**. Branched ^1^SGN-dendrites more commonly connected to AZs of neighboring IHCs (24 out of 29 of the 1^st^ sample and 42 out of 46 of the 2^nd^ sample) instead of to AZs of a single IHC. Branched ^1^SGNs preferentially contacted the modiolar face of the IHCs: 29 out of 29 (1^st^ sample) and 45 out of 46 (2^nd^ sample) contacted at least one modiolar AZ. Exclusive input from modiolar AZs was found for 13 out of 29 (1^st^ sample) and 23 out of 46 branched (2^nd^ sample) ^1^SGNs. As previously reported for the apex of the mouse cochlea^13^, AZs with multiple ribbons preferred the modiolar IHC side (20 out of 35, 1^st^ sample and 38 out of 48, 2^nd^ sample) also in our mid-cochlear datasets.

**Figure 3.**
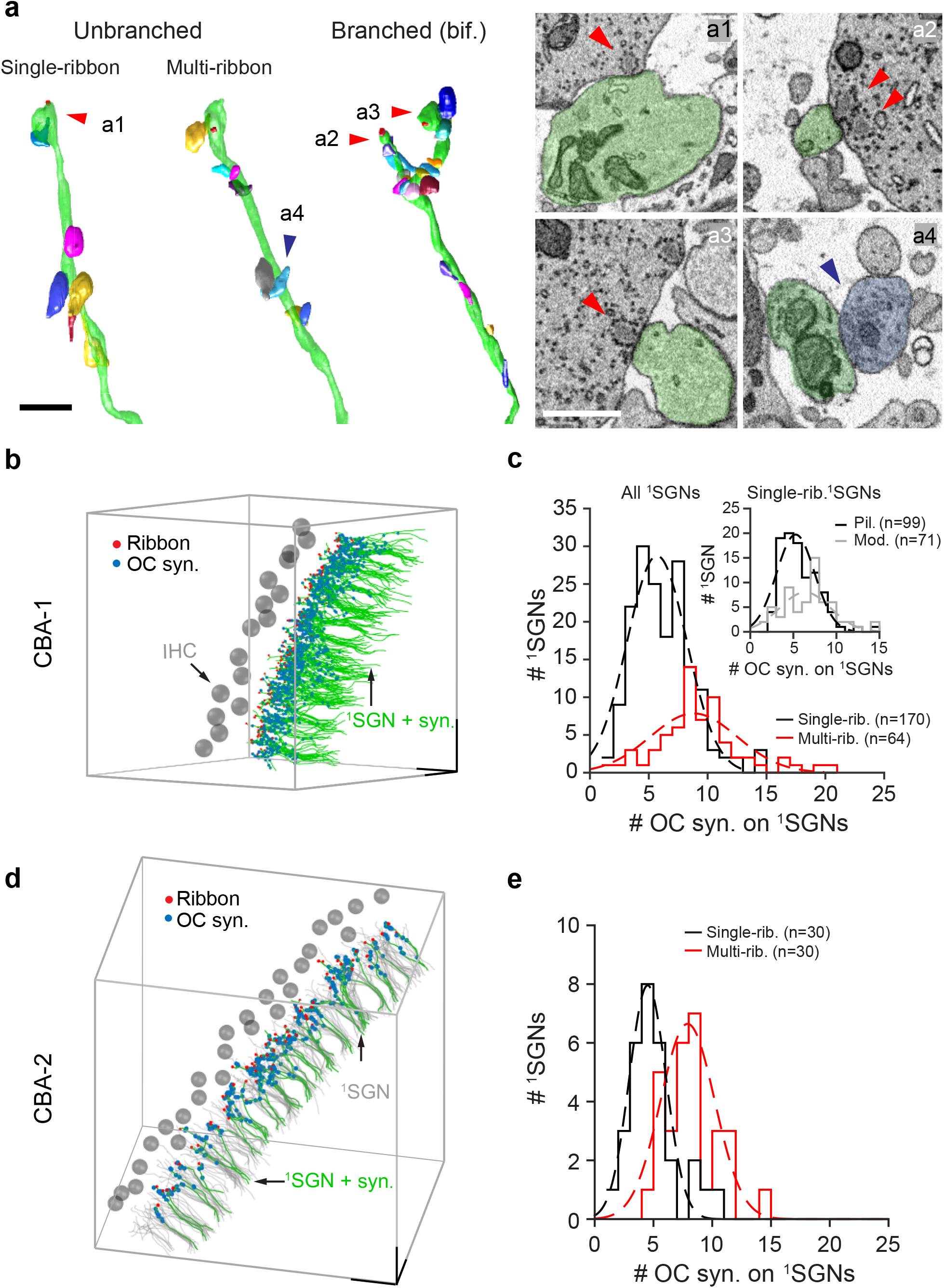
Quantification of afferent and efferent inputs on the ^1^SGN-dendrites. **(a)** Volume reconstruction of three distinct ^1^SGN classes characterized by single-ribbon (unbranched ^1^SGN, left), multi-ribbon (unbranched ^1^SGN, middle), and bifurcation (branched ^1^SGN, right). Scale bar 5 μm. Right: (**a1**-**a4**) cross-sections through synaptic structures indicated in (a) with arrows. Single-ribbon AZ (**a1**), dual-ribbon AZ (**a2**), ribbons on a bifurcated ^1^SGN-dendrite (**a2**& **a3**), and a representative efferent synapse on a ^1^SGN dendritic shaft (**a4**). Scale bar 1 μm**. (b)** Display of full-length unmyelinated ^1^SGN-dendrites (green) with presynaptic ribbons (red dots) and efferent synapses (OC inputs, blue dots) in the 1^st^ SBEM dataset. Scale bar 20 μm. **(c)** Histograms of the number of efferent synapses on ^1^SGN-dendrites grouped according to ribbon structures. Multi-ribbon ^1^SGNs (red, n = 64) received more efferent innervation than single-ribbon ^1^SGNs (black, n = 170), while the difference in efferent synapse number between ^1^SGNs postsynaptic to single modiolar ribbons (grey, n = 71) and those to single pillar ribbons (black, n = 99) was less prominent. On average 5.70 ± 2.53 and 8.82 ± 3.70 efferent synapses were found innervating ^1^SGN with single-(black) and multi-ribbon (red) synapses. For ^1^SGNs with single-ribbon synapses, the number of efferent synapses innervating pillar ^1^SGNs (black, inset) and modiolar ^1^SGNs (grey, inset) were 5.25 ± 2.19 and 6.47 ± 2.81, respectively. Two-sample t-tests suggested statistical significance between single- and multi-ribbon ^1^SGNs (***p <0.001) and between ^1^SGNs with single pillar and modiolar ribbon synapses (**p = 0.0028). (**d**) Display of full-length unmyelinated ^1^SGN-dendrites (grey, n = 266) in the 2^nd^ animal. 30 single- and 30 multi-ribbon ^1^SGNs (green) were randomly selected. Their presynaptic ribbons (red dots) and efferent synapses (OC inputs, blue dots) were annotated. Scale bar 20 μm. (**e**) Histograms of the number of efferent synapses on randomly selected ^1^SGN-dendrites grouped according to ribbon structures. On average 4.43 ± 2.16 (n = 30) and 7.70 ± 2.29 (n = 30) efferent synapses were found innervating ^1^SGNs with single-(black) and multi-ribbon (red) synapses. Two-sample t-test, ***p < 0.001.

Analysis of the efferent innervation of the unmyelinated segments of ^1^SGN-dendrites revealed a varying number of efferent synapses ranging from 0 up to 20 (**Fig. 3 b**). Significantly more efferent contacts are formed on unbranched and branched multi-ribbon ^1^SGNs than on unbranched, single-ribbon ^1^SGNs (**Fig. 3 c**. 8.82 ± 3.70 for multi-ribbon ^1^SGNs vs. 5.70 ± 2.53 for single-ribbon ^1^SGNs, p < 0.001). For unbranched, single-ribbon ^1^SGNs a stronger efferent innervation was observed for modiolar ^1^SGNs than for pillar ^1^SGNs (6.37 ± 2.95 for the modiolar ^1^SGNs vs. 5.26 ± 2.20 for the pillar ^1^SGNs, **Fig. 3 c inset,** p = 0.0073). Considering that AZs with large and multiple ribbons, which likely provide greater maximum rates of transmitter release, are more prevalent on the IHC modiolar side, our data indicates the strength of efferent modulation of ^1^SGNs correlates with stronger afferent input. We controlled this result by mapping all efferent innervation on randomly selected single- and multi-ribbon ^1^SGN-dendrites in the 2^nd^ animal (**Fig. 3 d**). Again, we found more efferent synapses formed on multi-ribbon ^1^SGN-dendrites than on those of single-ribbon ones (7.70 ± 2.29 vs. 4.43 ± 2.16, **Fig. 3 e**). Note that the increased efferent innervation was achieved by more efferent synapses along the ^1^SGN-dendrites in the ISB region rather than in the direct vicinity of ribbon synapse (**Suppl. Figure 3**).

### ^**1**^SGNs are innervated by efferent terminals of both LOC and MOC fibers

We note that the above analysis of efferent innervation did not distinguish synapses formed by LOC and MOC fibers. Due to the extent of the reconstruction, it was possible to distinguish MOC fibers from LOC fibers (**Fig. 4 a** to **c**), unambiguously by their characteristic crossing of the tunnel of Corti (**Fig. 4 d**) and OHC innervation (**Fig. 4 e**), while LOC fibers remain exclusively in the ISB region. Unexpectedly, we found that ^1^SGNs were also frequently contacted by MOC fibers, which showed parallel calibers of varying length with much less branching in the ISB before turning to cross the tunnel (**Fig. 4 d** to **f**). Compared to LOC synapses (**Fig. 4 c**), MOC terminals on ^1^SGN-dendrites showed larger bouton size but sparser vesicle content (**Fig. 4 f**). Extended structural quantification of all 39 MOC and 32 LOC fibers revealed that MOC fibers form fewer efferent synapses in the ISB region (**Fig. 4 g**) and preferentially innervate ^1^SGN-dendrites contacting the pillar IHC side (**Fig. 4 h**). This is consistent with the observation that MOC-innervated ^1^SGNs have small ribbon size (**Fig. 4 i**) and receive weaker efferent innervation (**Fig. 4 j**). In conclusion, this data suggests a hitherto unreported function of MOC fibers in modulating ^1^SGNs, particularly those contacting the IHC pillar side. Together with the preference of LOC fibers to innervate modiolar ^1^SGNs, the data indicates a differential modulation of ^1^SGNs by projections from medial and lateral subdivisions of the superior olivary complex.

**Figure 4.**
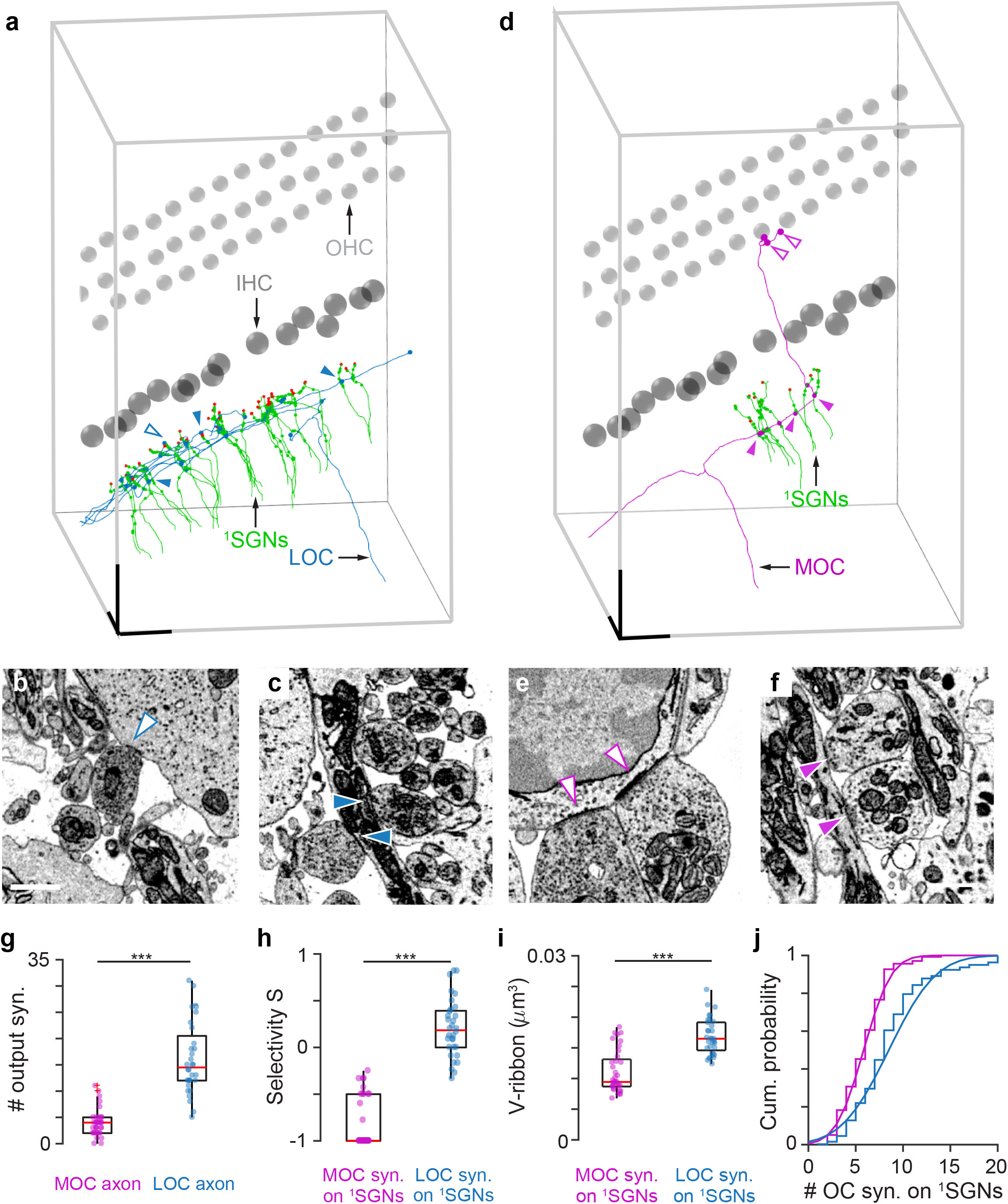
Efferent innervation of ^1^SGN-dendrites: selectivity of LOC and MOC control. **(a)** Display of a representative LOC fiber (blue) with all 25 innervated ^1^SGN-dendrites (green) in the dataset. Scale bar 20 μm. **(b-c)** High-resolution exemplary images of a presynaptic LOC terminal contacting an IHC (blue arrow, open) and LOC synapses with a ^1^SGN-dendrite (blue arrows, filled). Scale bar 1 μm. **(d)** Display of a representative MOC fiber (magenta) with all seven innervated ^1^SGN-dendrites (green). Scale bar 20 μm. **(e**& **f)** High-resolution example images of presynaptic MOC synapses onto an OHC (magenta arrow, open) and on a ^1^SGN-dendrite (magenta arrows, filled). Scale bar 1 μm. **(g)** Boxplot showing the comparison between the number of efferent synapses formed by individual MOC fibers (magenta) and LOC fibers (blue). **(h)** Innervation selectivity of MOC fibers (magenta) and LOC fibers (blue) on ^1^SGNs postsynaptic to IHC pillar versus modiolar face, selectivity index S = (#^1^SGN_modiolar_ – #^1^SGN_pillar_) / (# ^1^SGN_modiolar_ + #^1^SGN_pillar_). Note that 30 out of 39 MOC fibers exclusively innervated pillar ^1^SGNs. **(i)** Boxplot showing the mean ribbon size of MOC-innervated ^1^SGNs (magenta) compared to that of LOC-innervated ^1^SGNs (blue). **(j)** Cumulative probability distribution of the number of efferent synapses on individual MOC-innervated ^1^SGNs (magenta) compared to that of LOC-innervated ^1^SGNs (blue). (For g to i, two-sample t-test, ***p < 0.001; for j, two-sample Kolmogorov-Smirnov test, ***p < 0.001).

### Spatial organization of efferent innervation on ^1^SGNs

As described above, both LOC and MOC innervation coexists on individual ^1^SGN-dendrites and their presynaptic terminals have distinct appearances (**Fig. 4 c** & **f**). By these criteria, we identified 1175 putative LOC and 342 putative MOC terminals out of the total of 1517 efferent contacts on ^1^SGN-dendrites. Consistent with the relative innervation specificity of LOC and MOC projections reported above, the dendrite-based analysis of efferent input indicates that both modiolar ^1^SGNs, and multi-ribbon ^1^SGNs in particular, receive strong LOC innervations (**Fig. 5 a**) but weak MOC innervations (**Fig. 5 b**). In fact, MOC synapses skip more than half of modiolar and multi-ribbon ^1^SGNs (**Fig. 4 h**) and preferentially contact the pillar ^1^SGNs that in return featured only few LOC synapses. In conclusion, the efferent innervation showed a ^1^SGN-subdivision-specific dichotomy: LOC preferentially synapse on modiolar and multi-ribbon ^1^SGNs and MOC predominantly on pillar ^1^SGNs (**Fig. 5 c**).

**Figure 5.**
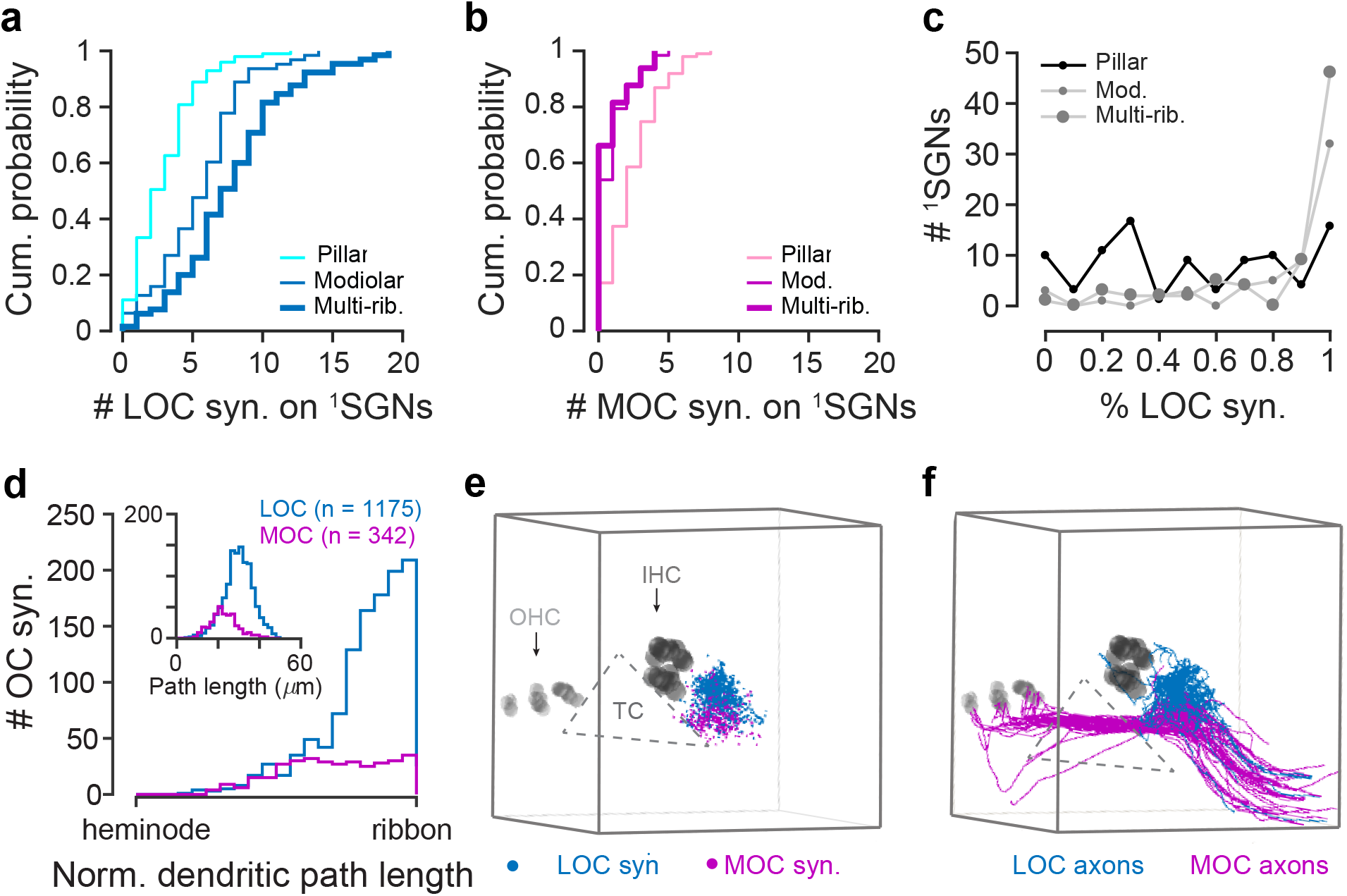
Spatial arrangement of OC innervation on ^1^SGN-dendrites. **(a)** Cumulative probability distribution of the number of LOC synapses on individual ^1^SGNs with pillar single-ribbon (light blue, thin line), modiolar single-ribbon (dark blue, thin line), and multi-ribbon (blue, thick line) contacts. **(b)** Similar to (a), cumulative probability distribution of the number of MOC synapses on individual ^1^SGNs with pillar single-ribbon (light magenta, thin line), modiolar single-ribbon (dark magenta, thin line), and multi-ribbon (magenta, thick line) contacts. Note that MOC synapses were lacking from more than 50 % modiolar and/or multi-ribbon ^1^SGNs (magenta). **(c)** Fraction of efferent synapses onto ^1^SGNs that are classified as LOC synapses with respect to pillar single-ribbon (small black dots), modiolar single-ribbon (small grey dots), as well as multi-ribbon (large grey dots) contacts. **(d)** Relative position of MOC synapses (magenta, n = 342) and LOC synapses (blue, n = 1175) on normalized dendritic paths between the ribbon-type AZ and heminode of ^1^SGN-dendrites. Inset: dendritic path lengths in μm measured from OC synapses to the heminode. **(e**& **f)**3D-illustration showing spatial segregation of LOC synapses (blue) from MOC synapses (magenta) (e), as well as traced trajectories of LOC (blue) from those of MOC fibers (magenta) (f). TC: Tunnel of Corti. (For a, b and d two-sample Kolmogorov-Smirnov test, ***p < 0.001).

Finally, we found a segregation of MOC and LOC innervation sites on ^1^SGN-dendrites with LOC terminals being more proximal to the ribbon synapses and MOC terminals synapsing more distally, towards the heminode (**Fig. 5 d inset**). We further measured the afferent dendritic pathlength and concluded that distinct innervation sites of LOC and MOC synapses were not a consequence of variability in their path length within the ISB (**Fig. 5 d**). A similar result was obtained from the same quantification on randomly selected ^1^SGNs in the 2^nd^ animal (**Suppl. Figure 4**). The spatial distribution of efferent terminals (**Fig. 5 e**) as well as the efferent fiber trajectories (**Fig. 5 f**) show a spatial segregation of the MOC and LOC projections within the ISB, which might instruct their preferred ^1^SGN innervation.

## Discussion

In the last decade, high throughput volume EM techniques have rendered dense reconstruction of large-scale mammalian neural circuits realistic within reasonable time and costs^58,59^. SBEM and FIB-SEM offer several advantages over other volume EM techniques based on thin slice manual collection, including less image distortion and slice loss, as well as fully automated acquisition cycle^59,60^. Despite lower z-resolution, in the end we chose SBEM instead of FIB-SEM in order to cover a larger volume for a more comprehensive circuit level analysis in the organ of Corti. Additional iterations of system optimization were made to overcome technical issues like charging artefacts and limited cutting thickness. Finally, by combining optimized sample preparation to increase tissue conductivity, implementing focal charge compensation to minimize charging from pure resin areas, and cutting along the direction of efferent calibers to improve the traceability at 40 nm z-resolution, we managed to acquire a dataset that allowed dense circuit reconstruction in the mature cochlea. The volume EM analysis of this study reveals an unprecedented complexity of the neural circuitry of the organ of Corti, rigorously tests established concepts on cochlear circuitry, and provides novel observations that allowed us to generate new hypotheses to be tested in the future. Novel findings regarding afferent and efferent innervation include a substantial fraction of branched SGN-dendrites and a differential efferent innervation of modiolar and pillar SGN-dendrites by LOC and MOC projections. Hence, afferent information mixing occurs and potentially improves the signal-to-noise relation of coding in a subset of ^1^SGNs. Efferent control of ^1^SGN-dendrites contacting the modiolar and pillar faces of the IHCs seems to involve fine-tuned innervation by MOC and LOC and likely contributes to diversifying ^1^SGN response properties for wide dynamic range sound encoding.

### Afferent connectivity of ^1^SGNs: graded synaptic strength and postsynaptic information mixing

The prevailing view on the afferent connectivity of ^1^SGNs in the mature mammalian cochlea is the mono-synaptic contact of an unbranched ^1^SGN-dendrite with a single-ribbon IHC AZ. Reconstructing a large volume of the mouse organ of Corti, we reveal a considerable fraction of branched ^1^SGN-dendrites (12 %). The bifurcation of the ^1^SGN-dendrite enables multiple synaptic contacts with one or, more frequently, two neighboring IHC(s) (**Fig. 3 a** and **Suppl. Figure 2**). Substantial branching of ^1^SGN-dendrites was reported for the human cochlea^51^, while its prevalence in other species such as guinea pig^49,61^ and cat^24,62^ remains less clear. We do not consider this to reflect a pathology as the number of afferent synapses per IHC (17.5 afferent synapses) agrees closely with other estimates for the mid-cochlear mouse organ of Corti obtained by immunofluorescence or electron microscopy^10,63,64^. The observation of branched, multi-ribbon ^1^SGNs adds further complexity to the afferent SGN connectivity: with regard to presynaptic IHCs, site of contact as well as ribbon morphology. It is tempting to speculate that such information mixing on the level of SGNs might improve the signal-to-noise ratio of sound coding. Peripheral branching of afferent auditory neurons is commonly found in the hearing organs of birds and lower vertebrates^65,66^.

So far, the presence of multiple ribbons at a single AZ was considered the exception to the rule and, if encountered, to occur at modiolar AZs^13,23,25,67^. Note that the multi-ribbon synapses described here and in other studies differ from complex synapses found in cat, which were defined as having an extended AZ with a single ribbon^24^. Here, we found that about 14 % afferent synapses contain more than one ribbon and together with another 12 % branched ^1^SGN with multiple afferent contacts constitute a considerable fraction (more than 1/4) of ^1^SGNs with enhanced excitation in the middle turn of the mouse cochlea (**Fig. 3 a**). In addition, we demonstrate the modiolar-pillar size gradient of ribbon size and abundance for IHC AZs at the level of electron microscopy for a large sample of 308 synapses (**Fig. 2 c**). This corroborates previous light microscopy^11,12^ and electron microscopy^23–25,67^ studies. Together with the notion of the preference of low spontaneous rate, high threshold ^1^SGNs for modiolar AZs, this begs the questions how presynaptic strength can relate to weaker sound responses. One candidate mechanism to explain this apparent conundrum is the more depolarized activation range of the Ca^2+^ influx at the modiolar AZs^12^. Our present study suggests that stronger efferent modulation of ^1^SGNs facing multi-ribbon AZs also contributes to ^1^SGN diversity (see next section).

As previously described for the mid-turn of the mouse cochlea by immunofluorescence microscopy^36^, we found the IHCs to be arranged in a partially staggered fashion. However, the influence of the staggered IHC arrangement on ribbon size seems minor if present at all (**Fig. 2**): IHCs closer to the pillar side tended to have smaller ribbons on average than those closer to the modiolus, which however did not reach significance. Interestingly, the notable negative correlation between the size and abundance of ribbons in IHCs (**Fig. 2 e**) suggests an endogenous control of total supply of ribbon material.

### Tuning the strength of afferent and efferent inputs into ^1^SGNs

A key result of the present study is the observation that the likely stronger maximal afferent input by multi-ribbon AZs and/or multiple synapses into SGNs is accompanied by stronger efferent innervation (**Fig. 3 c** & **e**). This contrasts findings in cat, where a modiolar-pillar gradient of efferent inputs on ^1^SGN-dendrites primarily resulted from efferent innervation of SGNs forming complex synapses at the IHCs’ modiolar face^24,25^. We postulate that the maximal synaptic strength of afferent transmission might co-determine the extent of efferent innervation by LOC projections. Previous studies showed a shifted acoustic sensitivity of ^1^SGNs to higher sound pressure levels during efferent activation^68,69^, resetting the dynamic range. We speculate that matching the strengths of afferent (ribbon) and efferent (LOC) transmission to modiolar ^1^SGNs may allow a tuning of sound encoding to fit the range of sound pressure “on demand”. Surgical de-efferentation of the mature mouse cochlea was shown to attenuate the modiolar-pillar gradient of ribbon size, suggesting a key role for the efferent system in maintaining functional heterogeneity of the afferent synapses^36^. Hence, efferent innervation might contribute to shaping neural response diversity underlying wide dynamic range coding by modulating ^1^SGNs as well as by instructing the synaptic properties in a position-dependent manner.

To our knowledge, this comprehensive EM circuit analysis of a large cochlear volume for the first time found MOC fibers to make considerable amounts of synaptic contacts with ^1^SGN-dendrites before leaving the ISB region towards OHCs (**Fig. 4 d**). Efferent ^1^SGN innervation seems to be a general feature of MOC innervation, because this kind of connections was found in 37 out of 39 annotated MOC fibers (**Fig. 4 g**), indicating a novel function of MOC modulation in ^1^SGN activities beyond its well-identified suppressive effect on OHC motility^32–34^. Previous EM studies, to our knowledge, did not report synapses onto ^1^SGN-dendrites in the ISB of tunnel-crossing MOC fibers, possibly because the reconstructed datasets were limited to 2-3 IHCs^13,24,25,51,53^. Moreover, as MOC fibers form *en passant* synapses instead of branched nerve endings with ^1^SGN-dendrites (**Fig. 4 d**), they likely escaped detection in single-fiber tracing experiments using light microscopy^52,70^. Nevertheless, *en passant* boutonal structures within the ISB can be appreciated in traced single MOC calibers^44^.

It remains to be studied whether MOC fibers employ acetylcholine as the only neurotransmitter in terminals on both ^1^SGN-dendrites and OHCs. If that was the case, based on the observation of an acetylcholine injection experiment^71^ one might expect an elevation of spontaneous firing in ^1^SGNs in addition to the classic MOC-mediated suppression of cochlear amplification. This way, MOC modulation might contribute to the high spontaneous firing rate of ^1^SGNs contacting the pillar IHC side (**Fig. 5 b**). MOC terminals tend to innervate SGNs closer to the heminode (**Fig. 5 d**), which may overlap with the proposed innervation site of putative inhibitory dopaminergic LOC fibers^72^. This notion seems supported by the observation in a LOC-lesion experiment that a narrow band of acetylcholine positive puncta colocalized partially with sparse dopamine positive puncta^45^. However, at least for the basal high frequency cochlea, such colocalization might alternatively reflect terminals of intrinsic cholinergic LOC neurons co-releasing dopamine^73^.

This study presents a comprehensive circuit diagram with quantification that enriches our insight into the structure underlying auditory signal processing. Besides novel insight into normal cochlear structure, the result of circuit analysis serves as a baseline for future structural investigations, including noise-induced synaptopathy^74^, aging-related structural alteration^75–77^ and putative tinnitus-related synaptic plasticity changes^78^.

## Acknowledgements

We thank Dr. M. Lei (SHIPM) for discussion and comments on the manuscript; Dr. M. Helmstaedter (MPI-BR) for support in the initial phase of the project; Drs. R. Redman, P. Bastians from Zeiss (APAC) for technical advice; Dr. J. Neef for comments on the manuscript; D. Li, J. Pan, J. Lu, H. Liu, B. Feng, Z. Jiang, C. Jin, L. Zhou, X. Yu, W. Wang, G. Li and Z. Chen for the tracing work. This study was supported by The Program for Professor of Special Appointment (Eastern Scholar) at Shanghai Institutions of Higher Learning (QD2018015 to Y.H.), the National Natural Science Foundation of China (81800901 to Y.H. and 81730028 to H.W.), the National Basic Research Development Program of China (SQ2017YFSF080012 to H.W.) and Shanghai Key Laboratory of Translational Medicine on Ear and Nose Diseases (14DZ2260300), Joint Research Initiative Shanghai Jiao Tong University School of Medicine (to Y.H. and T.M.). Work by T.M. was supported by the Deutsche Forschungsgemeinschaft (DFG, German Research Foundation) under Germany’s Excellence Strategy - EXC 2067/1-390729940 to T.M. as well as by the DFG’s Leibniz Program to T.M.

## Author contributions

Y.H., H.W. and T.M. designed and supervised the study. Y.H., X.D. and F.W. performed the SBEM experiment. Y.H. and H.H.W. analyzed the data. X.D., Y.G., Y.L. assisted with the data analysis. Y.H. and T.M. wrote the manuscript with the help of the other authors.

---

**Suppl. Figure 1. Quantification of tracing accuracy**(**a**) 7-fold skeletonization of the ^1^SGN-dendrites and an efferent axon traced by trained non-experts in the dataset. For ^1^SGN-dendrites, all tracing started from the ribbon-type AZ and proceeded to the heminode of the ^1^SGN-dendrite. Grey spheres represented the nuclei of IHCs which were arranged in a staggered manner. Scale bar 5 μm. (**b**-**d**) RESCOP analysis of traceability: histogram of edge votes (**b**), estimated edge-detection probability *p*(*p*_*e*_) distribution for all traced neurites (**c**), and prediction of tracing accuracy as a function of annotation redundancy (**d**).

**Suppl. Figure 2. Morphology of branched ^1^SGN-dendrites.**(**a**) Ten example skeletons of branched ^1^SGN-dendrites, on which ribbon synapses and OC synapses were indicated by red and blue dots, respectively. Scale bar 10 μm.

**Suppl. Figure 3. Similar efferent innervation of postsynaptic ^1^SGN boutons at the modiolar versus pillar IHC faces. (a)** Exemplary image of a ^1^SGN bouton (green). Efferent synapse (indicated by physical contact of SV-filled terminal) was marked by red asterisk and those within a 4-μm-sized box (white) but without physical contact by white asterisks. Scale bar 1 μm. **(b)** Boxplot of efferent synapse number on ^1^SGN boutons postsynaptic to IHC modiolar face (left) versus pillar face (right). Two-sample t test: p = 0.0837.

**Suppl. Figure 4. Same spatial arrangement of efferent innervation in the 2^nd^ animal**. (**a**) Dendritic pathlengths in μm measured from efferent synapses to the heminode. **(b)** Relative position of MOC synapses (magenta, n = 86) and LOC synapses (blue, n = 286) on normalized dendritic paths between ribbon AZ and heminode of ^1^SGN-dendrites.

## Material and Methods

### Animals

CBA/Ca mice were purchased from Sino-British SIPPR/BK Lab.Animal Ltd (Shanghai, China). This study was conducted at the Shanghai Institute of Precision Medicine and Ear Institute of Shanghai Ninth People’s Hospital. All procedures were reviewed and approved by the Institutional Authority for Laboratory Animal Care of the hospital (SH9H-2019-A387-1).

### Whole cochlea EM preparation

Animals were anesthetized through intraperitoneal injection of chloride hydrate (500 mg/kg) and temporal bones were removed after decapitation. The cochleae were fixed by perfusion through the round window with ice-cold fixative mixture containing 0.08 M cacodylate (pH 7.4), 2 % freshly-made paraformaldehyde (Sigma), and 2.5 % glutaraldehyde (Sigma), and then immersion-fixed for 5 hours, followed by a 4-hour decalcification in the same fixative with addition of 5 % EDTA (ethylenediaminetetraacetic acid, Sigma-Aldrich) at 4 °C.

The *en bloc* EM staining was performed following the previously published protocol^79^ with modifications. In brief, the decalcified cochleae were washed twice in 0.15 M cacodylate (pH 7.4) for 30 min each and sequentially immersed in 2 % OsO_4_ (Ted Pella), 2.5 % ferrocyanide (Sigma), and again 2 % OsO_4_ at room temperature for 2, 2, and 1.5 hours, respectively, without intermediate washing step. All staining solutions were buffered with 0.15 M cacodylate (pH 7.4). After being washed twice in nanopore filtered water for 30 min each, the cochleae were incubated at room temperature in filtered thiocarbonhydrazide (saturated aqueous solution, Sigma) for 1 hour, unbuffered OsO_4_ aqueous solution for 2 hours and lead aspartate solution (0.03 M, pH 5.0 adjusted by KOH, EMS) at 50 °C for 2 hours. Between steps, double rinses in nanopore filtered water for 30 min each were performed. For embedding, the cochleae were first dehydrated through a graded acetone series (50 %, 75 %, 90 %, 30 min each, all cooled at 4 °C) into pure acetone (3 × 100 %, 30 min at room temperature), followed by sequential infiltration with 1:1 and 1:2 mixtures of acetone and Spurr’s resin monomer (4.1 g ERL 4221, 0.95 g DER 736, 5.9 g NSA and 1 % DMAE; Sigma-Aldrich) at room temperature for 6 hours each on a rotator. Infiltrated cochleae were then incubated in pure resin overnight before being placed in embedding molds (Polyscience, Germany) and incubated in a pre-warmed oven (70 °C) for 72 hours.

### SBEM imaging of cochlea

Embedded samples were trimmed to a block-face of ~ 800 × 800 μm^2^ and imaged using a field-emission scanning EM (Gemini300, Zeiss) equipped with an in-chamber ultramicrotome (3ViewXP, Gatan) and back-scattered electron detector (Onpont, Gatan). For the 1^st^ CBA dataset, serial images were acquired in single tile mode (20,000 × 15,000 pixels) of 11 nm pixel size and nominal cutting thickness of 40 nm; incident beam energy 2 keV; dwell time 1 μs. 2500 slices were collected. For the 2^nd^ CBA dataset, 2952 serial images (16,000 × 9,000 pixels) were acquired at 12 nm pixel size and nominal cutting thickness of 50 nm; incident beam energy 2 eV; dwell time 1.5 μs. For both datasets focal charge compensation^80^ was set to 100 % with a high vacuum chamber pressure of ∼2.8 × 10^−3^ mbar. The datasets were aligned using self-written MATLAB script based on cross-correlation maximum between consecutive slices. Then the aligned datasets were split into cubes (128 × 128 × 128 voxels) for viewing and neurite-tracing in a browser-based annotation tool (webKNOSSOS ^56^).

### Neurite reconstruction and traceability test

Seed points were generated from 17 annotated afferent and one efferent terminal beneath an IHC at a central region of the dataset. These coordinates were delivered to annotators as starting points for neurite-tracing in all directions within the data volume. This yielded 19 × 7 = 133 skeletons with a total length of 7.185 mm, which were further analyzed using the RESCOP^57^ routine as described previously^79^. In brief, each set of seven redundant skeletons was compared computing the number of ‘pro’ votes and total votes for each edge in each skeleton-tracing. This resulted in a vote histogram that was corrected for the redundancy of each tracing, yielding the measured vote histogram. Next, the underlying prior of edge probability *p*(*p*_*e*_) was determined by fitting the vote histogram under the simplifying assumption that tracing decisions were independent between tracers and locations. Then, the predicted mean error-free path length as a function of the number of tracings per seed point (‘tracing redundancy’) was computed from the fitted prior *p*(*p*_*e*_) for the setting of annotators reconstructing a neurite.

### Ribbon size measurement and synapse counting

The ribbon size was measured by counting voxels which belonged to individual ribbon structures via volume-tracing tool in webKNOSSOS. Efferent synapse annotation was done by three independent annotators on traced skeletons and the result was proof-read by inspection of a 4^th^ annotator at annotated locations. All data analysis including statistic tests were conducted using self-written script and build-in functions in MATLAB (Mathworks).

## Additional information

**Suppl. Video 1:** Down-sampled cochlea SBEM data with alignment (scale bar 10 μm)

**Suppl. Video 2:** High resolution cochlea SBEM data in the ISB region (scale bar 2 μm)

## References

1 Moser, T., Grabner, C. P. & Schmitz, F. Sensory Processing at Ribbon Synapses in the Retina and the Cochlea. Physiol Rev 100, 103–144, doi:10.1152/physrev.00026.2018 (2020).

2 Fettiplace, R. Hair Cell Transduction, Tuning, and Synaptic Transmission in the Mammalian Cochlea. Compr Physiol 7, 1197–1227, doi:10.1002/cphy.c160049 (2017).

3 Huet, A. et al. Sound Coding in the Auditory Nerve: From Single Fiber Activity to Cochlear Mass Potentials in Gerbils. Neuroscience 407, 83–92, doi:10.1016/j.neuroscience.2018.10.010 (2019).

4 Reijntjes, D. O. J. & Pyott, S. J. The afferent signaling complex: Regulation of type I spiral ganglion neuron responses in the auditory periphery. Hear Res 336, 1–16, doi:10.1016/j.heares.2016.03.011 (2016).

5 Fuchs, P. A. & Lauer, A. M. Efferent Inhibition of the Cochlea. Cold Spring Harb Perspect Med 9, doi:10.1101/cshperspect.a033530 (2019).

6 Liberman, M. C. Single-neuron labeling in the cat auditory nerve. Science 216, 1239–1241, doi:10.1126/science.7079757 (1982).

7 Kiang, N. Y. S. Discharge patterns of single fibers in the cat’s auditory nerve. With the assistance of Takeshi Watanabe [et al.]. (M. I. T. Press, 1965).

8 Safieddine, S., El-Amraoui, A. & Petit, C. The auditory hair cell ribbon synapse: from assembly to function. Annu Rev Neurosci 35, 509–528, doi:10.1146/annurev-neuro-061010-113705 (2012).

9 Frank, T., Khimich, D., Neef, A. & Moser, T. Mechanisms contributing to synaptic Ca2+ signals and their heterogeneity in hair cells. Proc Natl Acad Sci U S A 106, 4483–4488, doi:10.1073/pnas.0813213106 (2009).

10 Meyer, A. C. et al. Tuning of synapse number, structure and function in the cochlea. Nat Neurosci 12, 444–453, doi:10.1038/nn.2293 (2009).

11 Liberman, L. D., Wang, H. & Liberman, M. C. Opposing gradients of ribbon size and AMPA receptor expression underlie sensitivity differences among cochlear-nerve/hair-cell synapses. J Neurosci 31, 801–808, doi:10.1523/JNEUROSCI.3389-10.2011 (2011).

12 Ohn, T. L. et al. Hair cells use active zones with different voltage dependence of Ca2+ influx to decompose sounds into complementary neural codes. Proc Natl Acad Sci U S A 113, E4716–4725, doi:10.1073/pnas.1605737113 (2016).

13 Michanski, S. et al. Mapping developmental maturation of inner hair cell ribbon synapses in the apical mouse cochlea. Proc Natl Acad Sci U S A 116, 6415–6424, doi:10.1073/pnas.1812029116 (2019).

14 Neef, J. et al. Quantitative optical nanophysiology of Ca(2+) signaling at inner hair cell active zones. Nat Commun 9, 290, doi:10.1038/s41467-017-02612-y (2018).

15 Liberman, M. C., Dodds, L. W. & Pierce, S. Afferent and efferent innervation of the cat cochlea: quantitative analysis with light and electron microscopy. J Comp Neurol 301, 443–460, doi:10.1002/cne.903010309 (1990).

16 Taberner, A. M. & Liberman, M. C. Response properties of single auditory nerve fibers in the mouse. J Neurophysiol 93, 557–569, doi:10.1152/jn.00574.2004 (2005).

17 Shrestha, B. R. et al. Sensory Neuron Diversity in the Inner Ear Is Shaped by Activity. Cell 174, 1229–1246 e1217, doi:10.1016/j.cell.2018.07.007 (2018).

18 Sun, S. et al. Hair Cell Mechanotransduction Regulates Spontaneous Activity and Spiral Ganglion Subtype Specification in the Auditory System. Cell 174, 1247–1263 e1215, doi:10.1016/j.cell.2018.07.008 (2018).

19 Petitpre, C. et al. Neuronal heterogeneity and stereotyped connectivity in the auditory afferent system. Nat Commun 9, 3691, doi:10.1038/s41467-018-06033-3 (2018).

20 Liberman, M. C. Auditory-nerve response from cats raised in a low-noise chamber. J Acoust Soc Am 63, 442–455, doi:10.1121/1.381736 (1978).

21 Winter, I. M., Robertson, D. & Yates, G. K. Diversity of characteristic frequency rate-intensity functions in guinea pig auditory nerve fibres. Hear Res 45, 191–202, doi:10.1016/0378-5955(90)90120-e (1990).

22 Sachs, M. B. & Abbas, P. J. Rate versus level functions for auditory-nerve fibers in cats: tone-burst stimuli. J Acoust Soc Am 56, 1835–1847, doi:10.1121/1.1903521 (1974).

23 Kantardzhieva, A., Liberman, M. C. & Sewell, W. F. Quantitative analysis of ribbons, vesicles, and cisterns at the cat inner hair cell synapse: correlations with spontaneous rate. J Comp Neurol 521, 3260–3271, doi:10.1002/cne.23345 (2013).

24 Liberman, M. C. Morphological differences among radial afferent fibers in the cat cochlea: an electron-microscopic study of serial sections. Hear Res 3, 45–63, doi:10.1016/0378-5955(80)90007-6 (1980).

25 Merchan-Perez, A. & Liberman, M. C. Ultrastructural differences among afferent synapses on cochlear hair cells: correlations with spontaneous discharge rate. J Comp Neurol 371, 208–221, doi:10.1002/(SICI)1096-9861(19960722)371:2<208::AID-CNE2>3.0.CO;2-6 (1996).

26 Furman, A. C., Kujawa, S. G. & Liberman, M. C. Noise-induced cochlear neuropathy is selective for fibers with low spontaneous rates. J Neurophysiol 110, 577–586, doi:10.1152/jn.00164.2013 (2013).

27 Song, Q. et al. Coding deficits in hidden hearing loss induced by noise: the nature and impacts. Sci Rep 6, 25200, doi:10.1038/srep25200 (2016).

28 Zhang, L., Engler, S., Koepcke, L., Steenken, F. & Koppl, C. Concurrent gradients of ribbon volume and AMPA-receptor patch volume in cochlear afferent synapses on gerbil inner hair cells. Hear Res 364, 81–89, doi:10.1016/j.heares.2018.03.028 (2018).

29 Heil, P. & Peterson, A. J. Basic response properties of auditory nerve fibers: a review. Cell Tissue Res 361, 129–158, doi:10.1007/s00441-015-2177-9 (2015).

30 Pangrsic, T., Singer, J. H. & Koschak, A. Voltage-Gated Calcium Channels: Key Players in Sensory Coding in the Retina and the Inner Ear. Physiol Rev 98, 2063–2096, doi:10.1152/physrev.00030.2017 (2018).

31 Warr, W. B. & Guinan, J. J., Jr. Efferent innervation of the organ of corti: two separate systems. Brain Res 173, 152–155, doi:10.1016/0006-8993(79)91104-1 (1979).

32 Evans, M. G., Lagostena, L., Darbon, P. & Mammano, F. Cholinergic control of membrane conductance and intracellular free Ca2+ in outer hair cells of the guinea pig cochlea. Cell Calcium 28, 195–203, doi:10.1054/ceca.2000.0145 (2000).

33 Dallos, P. et al. Acetylcholine, outer hair cell electromotility, and the cochlear amplifier. J Neurosci 17, 2212–2226 (1997).

34 Blanchet, C., Erostegui, C., Sugasawa, M. & Dulon, D. Acetylcholine-induced potassium current of guinea pig outer hair cells: its dependence on a calcium influx through nicotinic-like receptors. J Neurosci 16, 2574–2584 (1996).

35 Nouvian, R., Eybalin, M. & Puel, J. L. Cochlear efferents in developing adult and pathological conditions. Cell Tissue Res 361, 301–309, doi:10.1007/s00441-015-2158-z (2015).

36 Yin, Y., Liberman, L. D., Maison, S. F. & Liberman, M. C. Olivocochlear innervation maintains the normal modiolar-pillar and habenular-cuticular gradients in cochlear synaptic morphology. J Assoc Res Otolaryngol 15, 571–583, doi:10.1007/s10162-014-0462-z (2014).

37 Eybalin, M. Neurotransmitters and neuromodulators of the mammalian cochlea. Physiol Rev 73, 309–373, doi:10.1152/physrev.1993.73.2.309 (1993).

38 Ruel, J. et al. Physiology, pharmacology and plasticity at the inner hair cell synaptic complex. Hear Res 227, 19–27, doi:10.1016/j.heares.2006.08.017 (2007).

39 Darrow, K. N., Maison, S. F. & Liberman, M. C. Cochlear efferent feedback balances interaural sensitivity. Nat Neurosci 9, 1474–1476, doi:10.1038/nn1807 (2006).

40 Irving, S., Moore, D. R., Liberman, M. C. & Sumner, C. J. Olivocochlear efferent control in sound localization and experience-dependent learning. J Neurosci 31, 2493–2501, doi:10.1523/JNEUROSCI.2679-10.2011 (2011).

41 Eybalin, M., Caicedo, A., Renard, N., Ruel, J. & Puel, J. L. Transient Ca2+-permeable AMPA receptors in postnatal rat primary auditory neurons. Eur J Neurosci 20, 2981–2989, doi:10.1111/j.1460-9568.2004.03772.x (2004).

42 Ehret, G. & Frankenreiter, M. Quantitative analysis of cochlear structures in the house mouse in relation to mechanisms of acoustical information processing. Journal of Comparative Physiology 122, 66–85, doi:https://doi.org/10.1007/BF00611249 (1977).

43 Maison, S. F., Adams, J. C. & Liberman, M. C. Olivocochlear innervation in the mouse: immunocytochemical maps, crossed versus uncrossed contributions, and transmitter colocalization. J Comp Neurol 455, 406–416, doi:10.1002/cne.10490 (2003).

44 Warr, W. B. & Boche, J. E. Diversity of axonal ramifications belonging to single lateral and medial olivocochlear neurons. Exp Brain Res 153, 499–513, doi:10.1007/s00221-003-1682-3 (2003).

45 Darrow, K. N., Simons, E. J., Dodds, L. & Liberman, M. C. Dopaminergic innervation of the mouse inner ear: evidence for a separate cytochemical group of cochlear efferent fibers. J Comp Neurol 498, 403–414, doi:10.1002/cne.21050 (2006).

46 Fuchs, P. A., Lehar, M. & Hiel, H. Ultrastructure of cisternal synapses on outer hair cells of the mouse cochlea. J Comp Neurol 522, 717–729, doi:10.1002/cne.23478 (2014).

47 Thiers, F. A., Nadol, J. B., Jr. & Liberman, M. C. Reciprocal synapses between outer hair cells and their afferent terminals: evidence for a local neural network in the mammalian cochlea. J Assoc Res Otolaryngol 9, 477–489, doi:10.1007/s10162-008-0135-x (2008).

48 Liberman, M. C. Efferent synapses in the inner hair cell area of the cat cochlea: an electron microscopic study of serial sections. Hear Res 3, 189–204, doi:10.1016/0378-5955(80)90046-5 (1980).

49 Hashimoto, S., Kimura, R. S. & Takasaka, T. Computer-aided three-dimensional reconstruction of the inner hair cells and their nerve endings in the guinea pig cochlea. Acta Otolaryngol 109, 228–234, doi:10.3109/00016489009107438 (1990).

50 Nadol, J. B., Jr. Serial section reconstruction of the neural poles of hair cells in the human organ of Corti. II. outer hair cells. Laryngoscope 93, 780–791, doi:10.1288/00005537-198306000-00015 (1983).

51 Nadol, J. B., Jr. Serial section reconstruction of the neural poles of hair cells in the human organ of Corti. I. Inner hair cells. Laryngoscope 93, 599–614, doi:10.1002/lary.1983.93.5.599 (1983).

52 Brown, M. C. Morphology of labeled efferent fibers in the guinea pig cochlea. J Comp Neurol 260, 605–618, doi:10.1002/cne.902600412 (1987).

53 Brown, M. C. Morphology of labeled afferent fibers in the guinea pig cochlea. J Comp Neurol 260, 591–604, doi:10.1002/cne.902600411 (1987).

54 Denk, W. & Horstmann, H. Serial block-face scanning electron microscopy to reconstruct three-dimensional tissue nanostructure. PLoS Biol 2, e329, doi:10.1371/journal.pbio.0020329 (2004).

55 Willott, J. F. Handbook of mouse auditory research: from behavior to molecular biology. (CRC Press, 2001).

56 Boergens, K. M. et al. webKnossos: efficient online 3D data annotation for connectomics. Nat Methods 14, 691–694, doi:10.1038/nmeth.4331 (2017).

57 Helmstaedter, M., Briggman, K. L. & Denk, W. High-accuracy neurite reconstruction for high-throughput neuroanatomy. Nat Neurosci 14, 1081–1088, doi:10.1038/nn.2868 (2011).

58 Helmstaedter, M. Cellular-resolution connectomics: challenges of dense neural circuit reconstruction. Nat Methods 10, 501–507, doi:10.1038/nmeth.2476 (2013).

59 Kornfeld, J. & Denk, W. Progress and remaining challenges in high-throughput volume electron microscopy. Curr Opin Neurobiol 50, 261–267, doi:10.1016/j.conb.2018.04.030 (2018).

60 Briggman, K. L. & Bock, D. D. Volume electron microscopy for neuronal circuit reconstruction. Curr Opin Neurobiol 22, 154–161, doi:10.1016/j.conb.2011.10.022 (2012).

61 Fernandez, C. The innervation of the cochlea (guinea pig). Laryngoscope 61, 1152–1172, doi:10.1288/00005537-195112000-00002 (1951).

62 Spoendlin, H. Innervation patterns in the organ of corti of the cat. Acta Otolaryngol 67, 239–254, doi:10.3109/00016486909125448 (1969).

63 Stamataki, S., Francis, H. W., Lehar, M., May, B. J. & Ryugo, D. K. Synaptic alterations at inner hair cells precede spiral ganglion cell loss in aging C57BL/6J mice. Hear Res 221, 104–118, doi:10.1016/j.heares.2006.07.014 (2006).

64 Buran, B. N. et al. Onset coding is degraded in auditory nerve fibers from mutant mice lacking synaptic ribbons. J Neurosci 30, 7587–7597, doi:10.1523/JNEUROSCI.0389-10.2010 (2010).

65 Fischer, F. P. Quantitative analysis of the innervation of the chicken basilar papilla. Hear Res 61, 167–178, doi:10.1016/0378-5955(92)90048-r (1992).

66 Klinke, R. Neurotransmission in the inner ear. Hear Res 22, 235–243, doi:10.1016/0378-5955(86)90100-0 (1986).

67 Jean, P. et al. The synaptic ribbon is critical for sound encoding at high rates and with temporal precision. Elife 7, doi:ARTN e29275 10.7554/eLife.29275 (2018).

68 Guinan, J. J., Jr. & Gifford, M. L. Effects of electrical stimulation of efferent olivocochlear neurons on cat auditory-nerve fibers. I. Rate-level functions. Hear Res 33, 97–113, doi:10.1016/0378-5955(88)90023-8 (1988).

69 Winslow, R. L. & Sachs, M. B. Effect of electrical stimulation of the crossed olivocochlear bundle on auditory nerve response to tones in noise. J Neurophysiol 57, 1002–1021, doi:10.1152/jn.1987.57.4.1002 (1987).

70 Liberman, M. C. & Brown, M. C. Physiology and anatomy of single olivocochlear neurons in the cat. Hear Res 24, 17–36, doi:10.1016/0378-5955(86)90003-1 (1986).

71 Felix, D. & Ehrenberger, K. The efferent modulation of mammalian inner hair cell afferents. Hear Res 64, 1–5, doi:10.1016/0378-5955(92)90163-h (1992).

72 Maison, S. F. et al. Dopaminergic signaling in the cochlea: receptor expression patterns and deletion phenotypes. J Neurosci 32, 344–355, doi:10.1523/JNEUROSCI.4720-11.2012 (2012).

73 Wu, J. S. et al. Sound exposure dynamically induces dopamine synthesis in cholinergic LOC efferents for feedback to auditory nerve fibers. Elife 9, doi:10.7554/eLife.52419 (2020).

74 Kujawa, S. G. & Liberman, M. C. Adding insult to injury: cochlear nerve degeneration after “temporary” noise-induced hearing loss. J Neurosci 29, 14077–14085, doi:10.1523/JNEUROSCI.2845-09.2009 (2009).

75 Lauer, A. M., Fuchs, P. A., Ryugo, D. K. & Francis, H. W. Efferent synapses return to inner hair cells in the aging cochlea. Neurobiol Aging 33, 2892–2902, doi:10.1016/j.neurobiolaging.2012.02.007 (2012).

76 Fernandez, K. A., Jeffers, P. W., Lall, K., Liberman, M. C. & Kujawa, S. G. Aging after noise exposure: acceleration of cochlear synaptopathy in “recovered” ears. J Neurosci 35, 7509–7520, doi:10.1523/JNEUROSCI.5138-14.2015 (2015).

77 Sergeyenko, Y., Lall, K., Liberman, M. C. & Kujawa, S. G. Age-related cochlear synaptopathy: an early-onset contributor to auditory functional decline. J Neurosci 33, 13686–13694, doi:10.1523/JNEUROSCI.1783-13.2013 (2013).

78 Knudson, I. M., Shera, C. A. & Melcher, J. R. Increased contralateral suppression of otoacoustic emissions indicates a hyperresponsive medial olivocochlear system in humans with tinnitus and hyperacusis. J Neurophysiol 112, 3197–3208, doi:10.1152/jn.00576.2014 (2014).

79 Hua, Y., Laserstein, P. & Helmstaedter, M. Large-volume en-bloc staining for electron microscopy-based connectomics. Nat Commun 6, 7923, doi:10.1038/ncomms8923 (2015).

80 Deerinck, T. J. et al. High-performance serial block-face SEM of nonconductive biological samples enabled by focal gas injection-based charge compensation. J Microsc 270, 142–149, doi:10.1111/jmi.12667 (2018).

