## Supplementary figures and images for "Electron Microscopic Reconstruction of Neural Circuitry in the Cochlea"

### Supplemental Figures

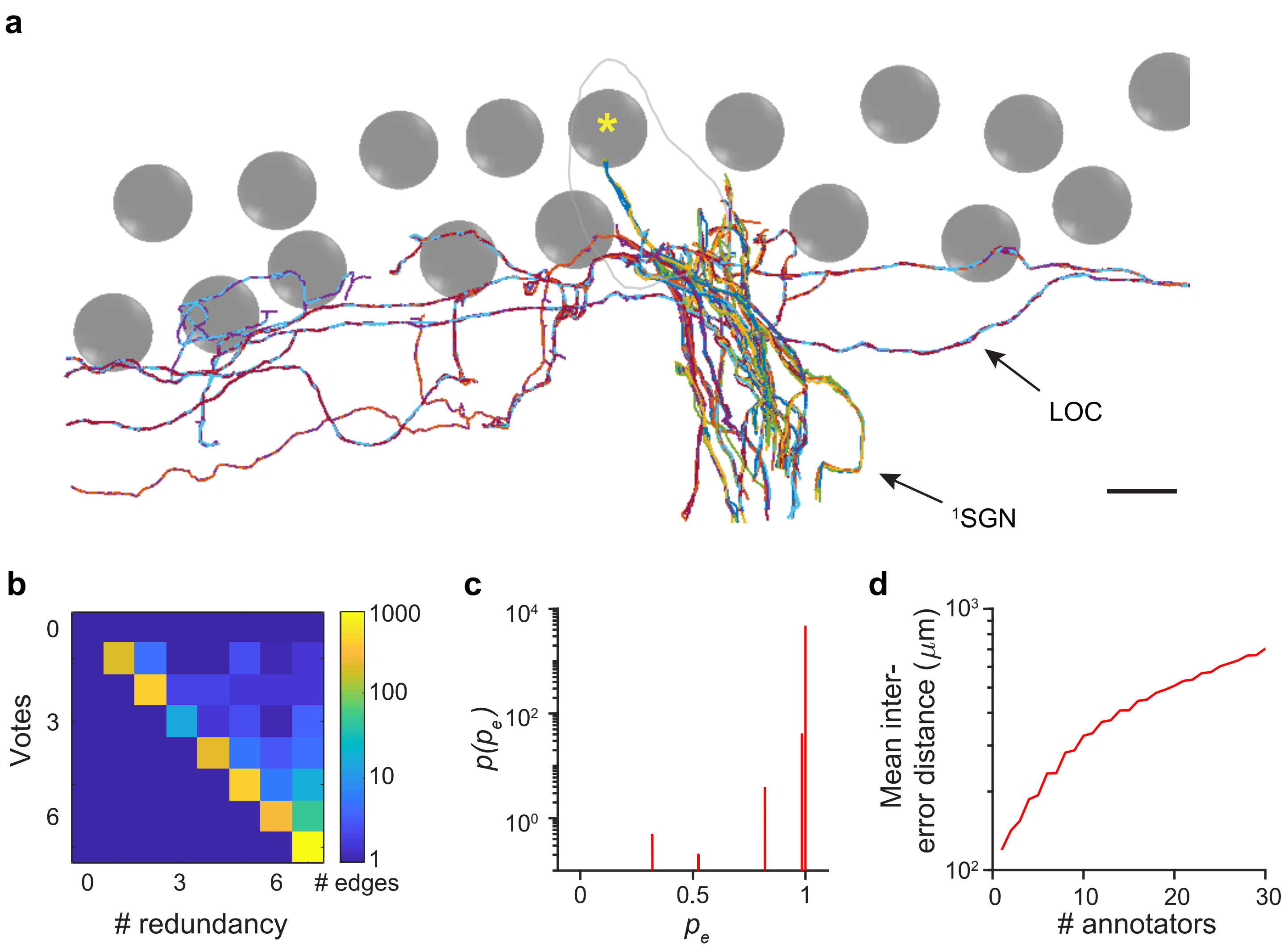

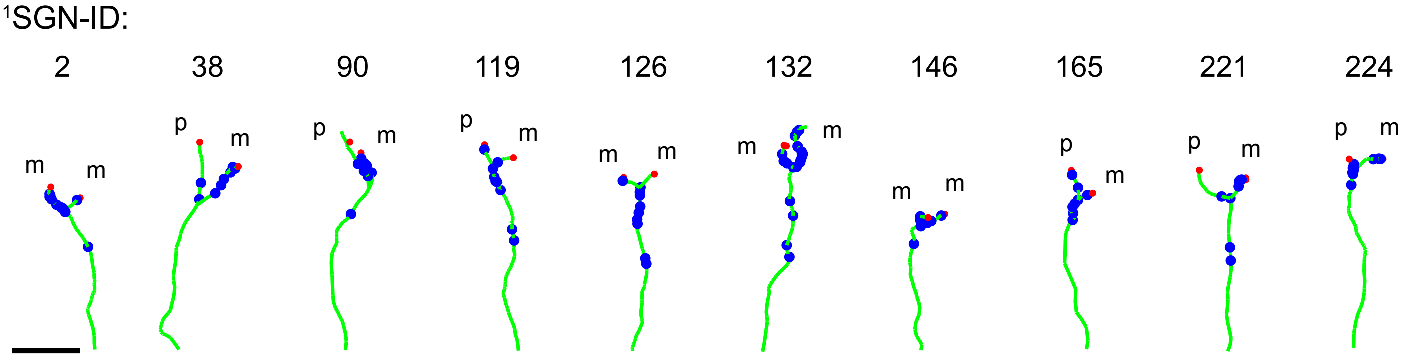

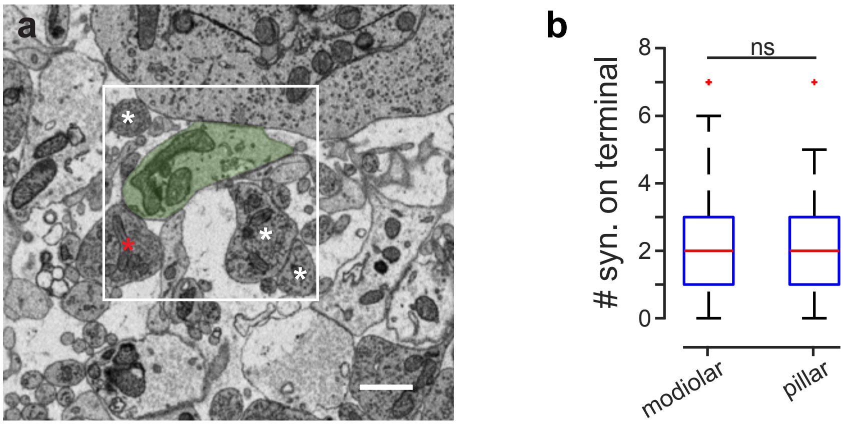


**Suppl. 3**

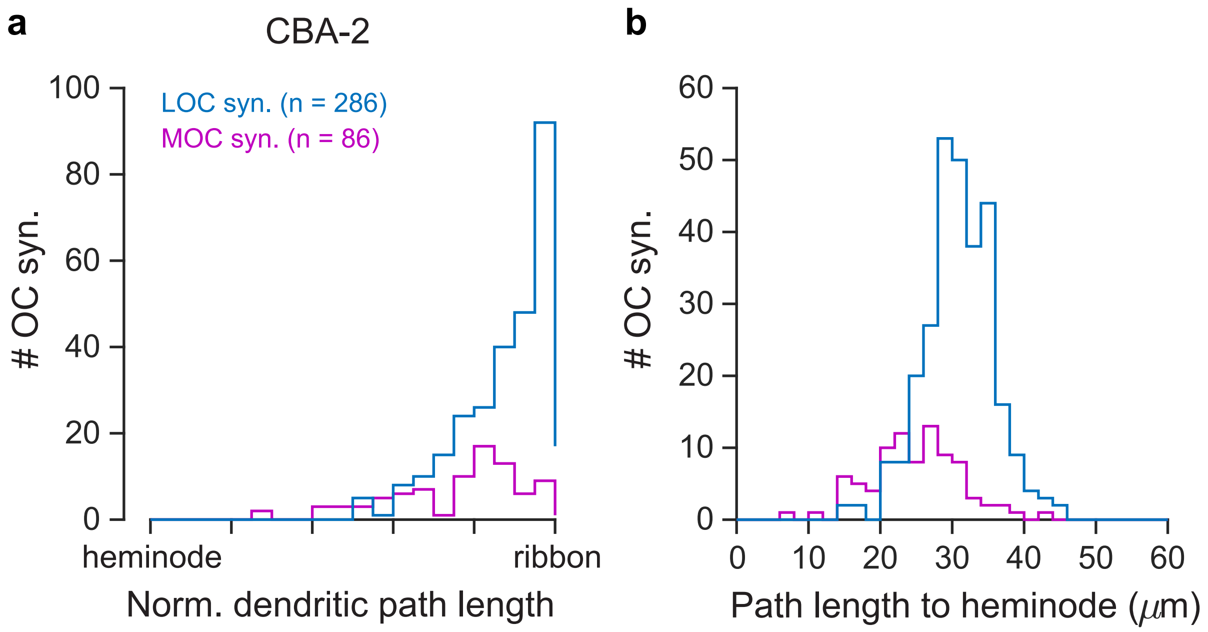
